# Conifer leaves have a peroxisomal oxidative decarboxylation path in the photorespiratory pathway

**DOI:** 10.1101/2022.02.11.480092

**Authors:** Shin-Ichi Miyazawa, Takafumi Miyama, Ko Tahara, Tokuko Ujino-Ihara, Hiroyuki Tobita, Yuji Suzuki, Mitsuru Nishiguchi

## Abstract

The photorespiratory pathway consists of enzymes operating in chloroplasts, mitochondria, and peroxisomes. Conifer leaves lack one of them, chloroplastic Gln synthetase, which questioned the current belief that the photorespiratory mechanism is identical between angiosperm C_3_ species and conifers. A photorespiratory-metabolite analysis of the leaves of 13 conifer and 14 angiosperm tree species revealed significant differences in the mean metabolite concentrations between the two taxonomic groups: the glycerate content on chlorophyll basis in conifer leaves was <1/10 that detected in angiosperm leaves, whereas the glycolate content was 1.6 times higher in conifer leaves. Glycerate is produced from Ser through an intermediate, hydroxypyruvate. To investigate the lower glycerate levels observed in conifer leaves, we performed experiments of ^13^C-labeled Ser feeding to the detached shoots of a conifer (*Cryptomeria japonica*) via the transpiration stream, and compared the labeling patterns of photorespiratory metabolites with those of an angiosperm (*Populus nigra*). Glycerate was most labeled in *P. nigra*, whereas glycolate was more labeled than glycerate in *C. japonica*. The photorespiration pathway involves H_2_O_2_-scavenging and H_2_O_2_-generating enzymes, catalase (CAT) and glycolate oxidase (GLO), respectively, which are the peroxisomal targeting enzymes in angiosperms. In contrast, database analyses of the peroxisomal targeting signal motifs and analyses of the peroxisomal fractions isolated from *C. japonica* leaves indicated that the conifer peroxisomes were not a major localization of CAT. These results suggest that the conifer photorespiration pathway has a bypass from Ser to glycolate via the decarboxylation of hydroxypyruvate, because of an imbalance between CAT and GLO activities in peroxisomes.

**One sentence summary:** Conifer peroxisome is not a major localization of catalase and yields a unique oxidative decarboxylation path in the photorespiratory pathway.

## Introduction

Photorespiration starts with the oxygenation of ribulose-1,5-bisphosphate (RuBP) by the CO_2_-fixation enzyme Rubisco. The oxygenation reaction with one molecule of RuBP produces one molecule each of 2-phosphoglycolate (2-PG) and the Calvin–Benson cycle intermediate 3-phosphoglycerate (3-PGA). 2-PG is a toxic product in plants and is converted to 3-PGA through the photorespiratory pathway, requiring not only CO2 emission, but also energy (ATP and reductants such as NADH and reduced ferredoxin) consumption. The pathway consists of at least 15 enzymes, including chloroplastic, mitochondrial, and peroxisomal enzymes, as well as transport proteins to transfer the intermediates between the cell compartments (Reumann and Weber, 2006; Reumann et al., 2007). Studies using *Arabidopsis* and barley mutants contributed to the clarification of the enzymes that participate in the photorespiratory pathway (Ogren, 1984; Somerville, 2001).

The glutamine synthetase (GS) located in chloroplasts, GS2, is an essential enzyme of the photorespiratory pathway (Wallsgrove et al., 1987). GS2 assimilates the ammonia (NH_3_) that is produced during the Gly-to-Ser conversion mediated by glycine decarboxylase complex (GDC) reactions in mitochondria. Conversely, GS2 is absent from the leaves of conifers, and it is considered that one of the cytosolic types of GS could be a substitute (Suárez et al., 2002; Cánovas et al., 2007; Miyazawa et al., 2018). The leaves of conifers such as *Cryptomeria japonica* and *Pinus densiflora* exhibited higher NH_3_ emission potentials than did angiosperm leaves, indicating a lowered NH3 assimilation rate during photorespiration in conifers (Miyazawa et al., 2018). These findings question our current belief that the photorespiratory mechanism of conifer leaves is identical to that of angiosperm species.

The rate of photorespiratory CO_2_ release concomitantly with the production of NH_3_ by the GDC reactions affects the net CO_2_ assimilation rates (*A*) of the leaves. The photosynthesis model (Farquhar–von Caemmerer–Berry model [FvCB model]) describes *A* via the following equation (Farquhar et al., 1980):

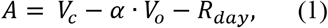

where *V_c_* and *V_o_* denote the carboxylation and oxygenation rates of Rubisco, respectively; And α and *R*_day_ are the rate of photorespiratory CO_2_ release per unit *V_o_* and the mitochondrial respiration rate in the light, respectively. By introducing the chloroplastic CO_2_ compensation points (*Γ*;* Farquhar et al. 1980), *A* is then given by:

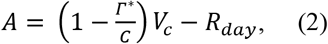

where

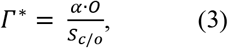

where *C* and *O* are the CO_2_ and O_2_ concentrations in chloroplasts, respectively; and *S_c/o_* denotes the Rubisco specificity factor. The *α* value is frequently assumed to be 0.5 because one molecule of CO_2_ during the GDC reaction is released per every two molecules of 2-PG (Farquhar et al., 1980).

The *Γ*\* value is crucial because it determines the rate of photorespiratory CO_2_ release at a given *C*. When *α* is set to 0.5 and *O* is constant, the *Γ*\* values are calculated from *in vitro* determinations of the *S*_c/o_ of Rubisco purified from leaves, which have been intensively reported for crop species (Hermida-Carrera et al., 2016). *Γ*\* values in tree species have rarely been reported, perhaps because of the difficulties in determining *S*_c/o_ in vitro. Conversely, without direct measurements of *S*_c/o_ in vitro and any assumptions of *α*, our previous study demonstrated that *Γ*\* could be estimated via an in vivo O_2_ response analysis of leaf CO_2_ compensation points (*Γ*, the CO_2_ concentrations in the air surrounding the leaves when *A* equals zero), and that the estimated *Γ*\* values of conifers such as *C. japonica* and *Metasequoia glyptostroboides* using this in vivo method did not significantly differ from those of herbaceous species, such as tobacco and French bean (Miyazawa et al., 2020). However, a similar *Γ*\* does not ensure that conifers have the same photorespiratory mechanism as angiosperms, because the efflux of CO_2_ escaping from inside the leaves via the photorespiratory pathway can be less than 0.5 or more in some instances (Cousins et al., 2008, 2011; Busch et al., 2013, 2020).

To address the question of whether conifer leaves have a different pathway from the orthodox photorespiratory pathway, first, we compared the photorespiratory-metabolite concentrations in leaves between various conifer and angiosperm tree species grown in a field site. Furthermore, we conducted experiments using the saplings of an angiosperm tree (*Populus nigra*) and a conifer (*C. japonica*), which were both cultivated in a growth chamber—gene information for the two species is provided in our public cDNA library (Forest EST and Genome database: ForestGEN)—to unravel the physiological mechanisms that are responsible for the differences in the photorespiratory-metabolite profiles detected between conifer and angiosperm tree leaves.

## Results

### Photorespiratory metabolites

We compared the concentrations of the photorespiratory metabolites of leaves between 13 conifer and 14 angiosperm tree species (Figure 1; Supplemental Table S1). The concentrations of all metabolites were expressed per unit leaf chlorophyll concentration. The metabolite concentrations were analyzed using the leaf samples collected in two consecutive years. The observed trends in the differences between conifers and angiosperm trees were unaffected by their sampling years. The results presented in Figure 1 were summarized on a map of the orthodox photorespiratory pathway (Figure 2).

**Figure 1.**
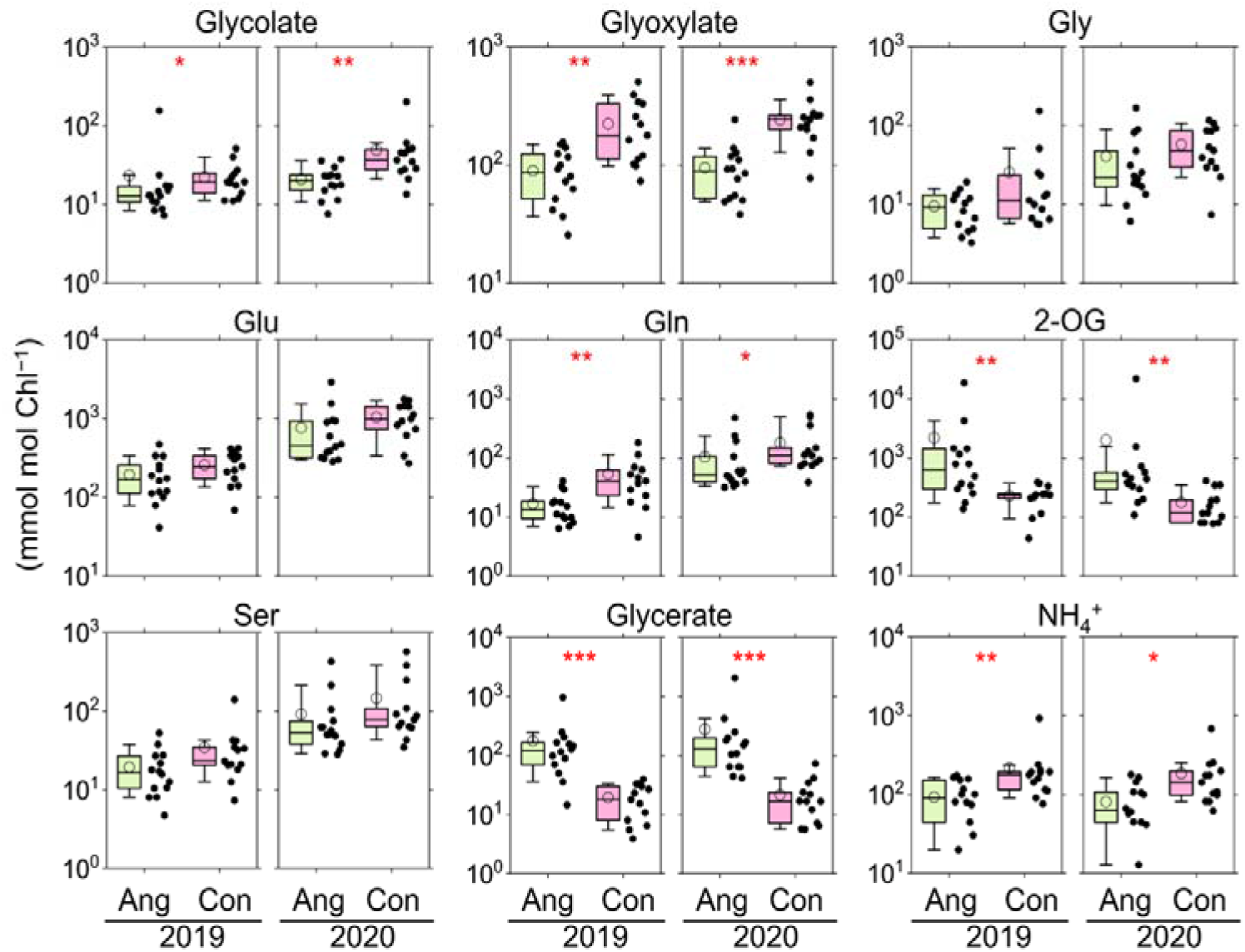
Comparison of leaf photorespiratory-metabolite concentrations on a chlorophyll basis between 14 angiosperm tree (Ang) and 13 conifer (Con) species grown in a field site. The left and right panels of each metabolite represent the samples collected in 2019 and 2020, respectively. The central box in each box plot represents the interquartile range and median; the whiskers indicate the 10^th^ and 90^th^ percentiles. The open circles indicate the average values. The scatter plots represent the data distributions. Mann Whitney *U* tests were conducted for the pairwise comparisons between angiosperms and conifers: *, *P* < 0.05; **, *P* < 0.01; ***, *P* < 0.001.

**Figure 2.**
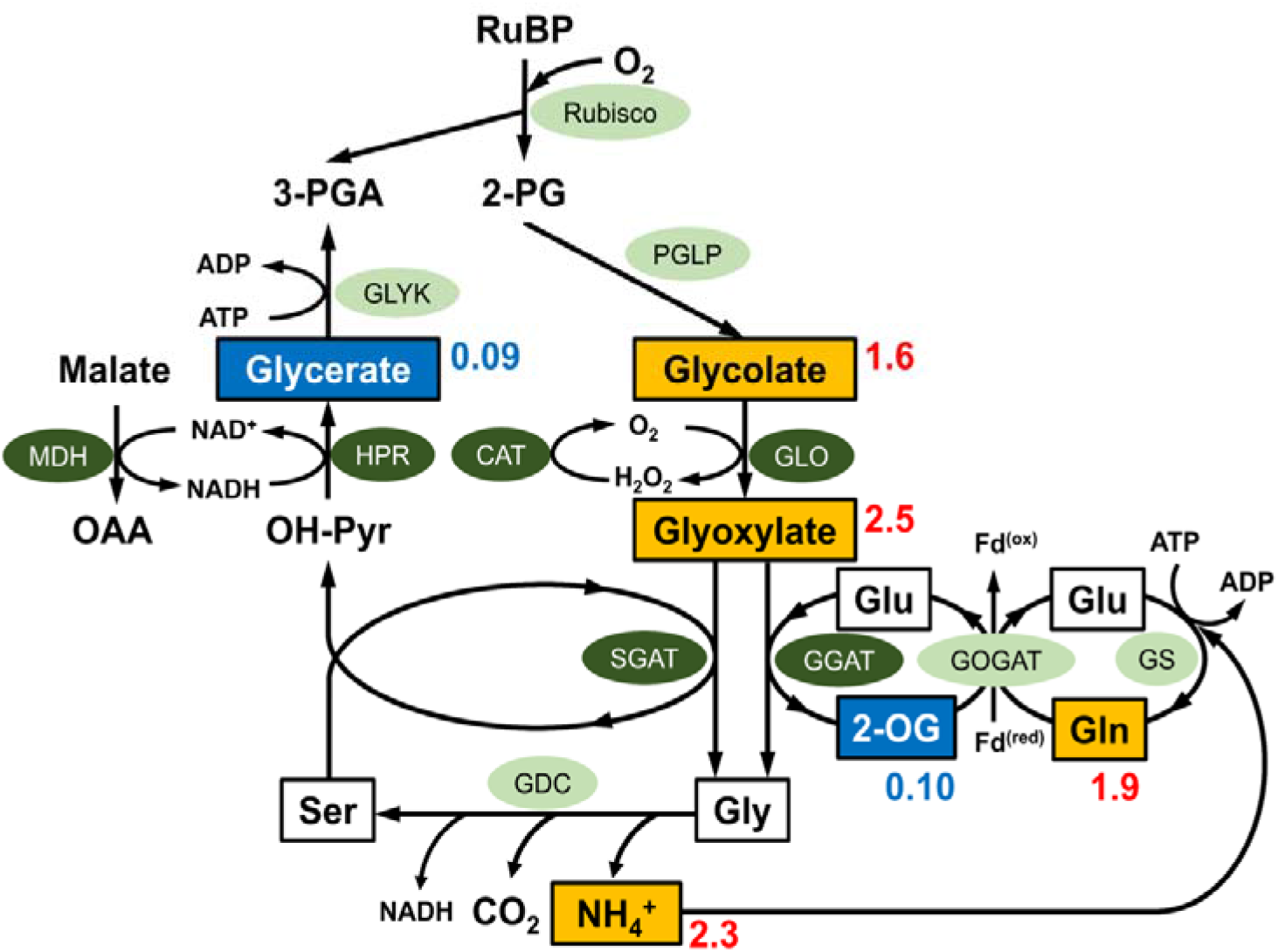
Difference in the metabolite concentrations of the orthodox photorespiratory pathway between the leaves of angiosperm trees and those of conifers. The values adjacent to each metabolite species represent the ratio of the two-year average concentration of the 13 conifer species to that of the 14 angiosperm tree species (Supplemental Table S1). The measured metabolites are boxed, and those that showed statistically significant differences between the two taxonomic groups are shown in color based on the calculated ratios (blue, decrease; orange, increase). Fd^(ox)^, oxidized ferredoxin; Fd^(red)^, reduced ferredoxin; 2-OG, 2-oxoglutarate; OΛA, oxaloacetate; OH-Pyr, hydroxypyruvate; 2-PG, 2-phosphoglycolatc; 3-PGA, 3-phosphoglyccratc; RuBP, ribulose-1,5-bisphosphatc. The enzymes arc indicated by circles (peroxisome-targeting enzymes are denoted in white letters in dark-green circles): CAT, catalase; GLO, glycolate oxidase; GDC, glycine decarboxylase complex; GGAT, glutamate-glyoxylatc aminotransferase; GLYK, glycerate kinase; GOGAT, glutamatc-2-OG aminotransferase; GS, glutamine synthetase; HPR, hydroxypyruvate reductase; MDH, malate dehydrogenase; PGLP, phosphoglycolate phosphatase; SGAT, scrine-glyoxylatc aminotransferase.

We found clear differences in the concentrations of several metabolite between the two taxonomic groups: the mean concentration of glycerate in conifer leaves was less than one-tenth that detected in angiosperm leaves (Figure 2). In contrast, the glycolate and glyoxylate concentrations in conifer leaves were 1.6 and 2.5 times higher than those observed in angiosperm leaves, respectively. The concentrations of 2-oxoglurtarate (2-OG), Gln, and NH_4_^+^ in the GS–GOGAT cycle also differed: the concentration of 2-OG in conifer leaves was about one-tenth that of angiosperm tree leaves, whereas those of Gln and NH_4_^+^ in conifer leaves were about twice higher than those recorded in angiosperm tree leaves. The concentrations of amino acids such as Gly, Ser, and Glu did not differ significantly between the two groups. Significantly lower levels of glycerate and 2-OG were also detected in conifer leaves when the metabolite concentrations were expressed per unit leaf fresh weight (Supplemental Figure S1).

### ^13^C-Ser labeling

In the orthodox photorespiration pathway, glycerate is produced from Ser through the sequential reactions of Ser-glyoxylate aminotransferase (SGAT) and hydroxypyruvate reductase (HPR) (Figure 2). To examine the mechanisms underlying the lower levels of glycerate detected in conifer leaves, we carried out a ^13^C-labeled Ser (^13^C-Ser) feeding experiment using detached shoots of a conifer, *C. japonica*. We examined which photorespiratory metabolite was labeled with ^13^C using gas chromatography-mass spectrometry (GC-MS) and compared the results with the labeling patterns of an angiosperm tree, *P. nigra*.

As inferred from the pathway, glycerate was mainly labeled with ^13^C in *P. nigra* (Figure 3; Supplemental Table S2). Conversely, the most labeled metabolite in *C. japonica* was glycolate, although glycerate was also labeled in this species: glycolate was 2–4.5 times more labeled than glycerate. In both tree species, other photorespiratory metabolites, such as 2-OG, Glu, and Gln, were not significantly labeled, with the exception of Gly. Thus, the metabolite labeling pattern was different between *P. nigra* and *C. japonica*.

**Figure 3.**
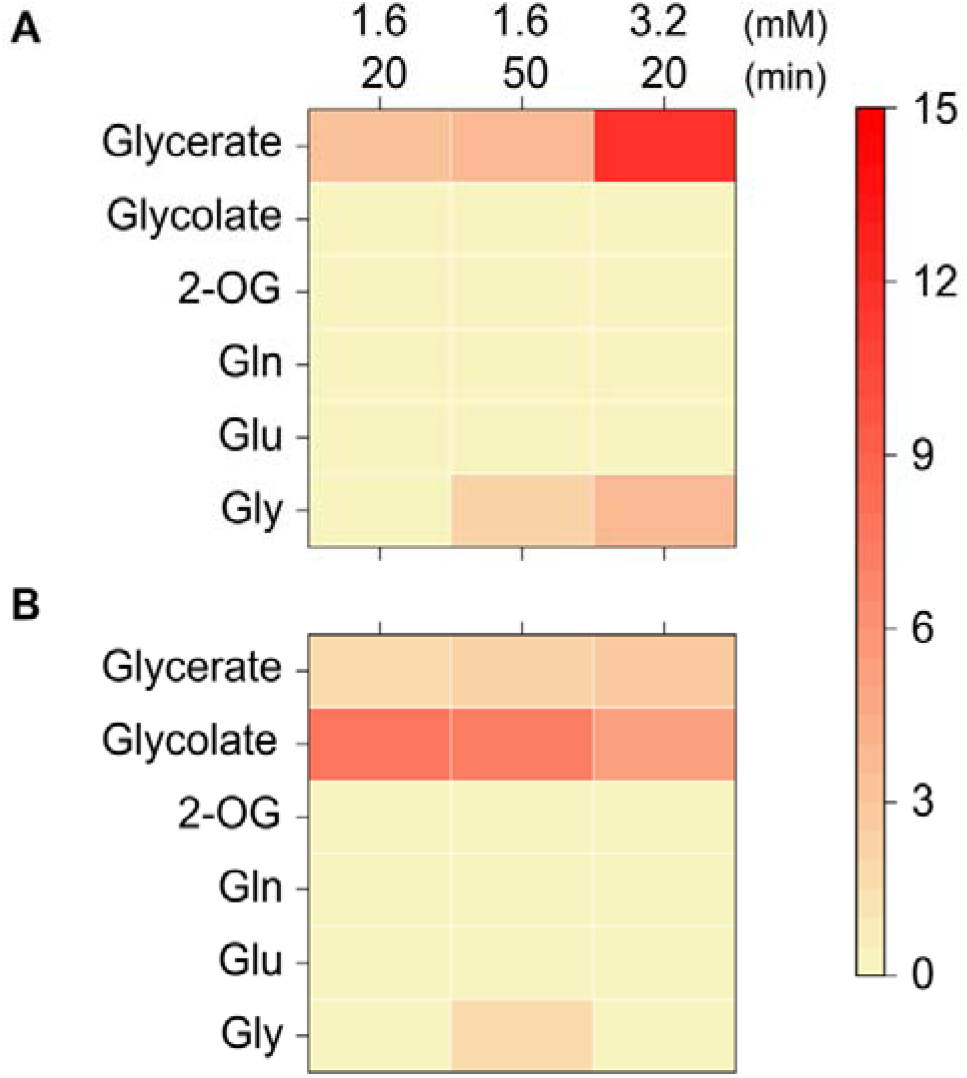
Heatmap visualization of the leaf photorespiratory metabolites labeled with a stable isotope of carbon (^13^C). Ser (1.6 or 3.2 mM) was prepared so that its C3 position was labeled with ^13^C (^13^C-Ser) and was then fed to the detached shoots of an angiosperm tree (*Populus nigra*) (A) and those of a conifer (*Cryptomería japonica*) (B) via the transpiration stream. As a control, unlabclcd standard Scr (uScr) feedings were also carried out. The feedings were conducted over 20 or 50 min under illumination in an environmentally controlled growth cabinet. The row displays the metabolite species; the column displays different treatments. The brightness of the red color coσcsponds to the magnitude of the increases in the relative abundance of the isotopologue, which had a single ^13^C atom for each metabolite (Student’s *t*-test; *P* < 0.05 between the ^13^C-Scr and uScr feedings); the yellow color indicates the absence of significant differences. The number of shoot samples used in each treatment per species was 3–4 (Supplemental Table S2).

### Enzyme activities

Glycerate is produced from hydroxypyruvate (OH-Pyr) by HPR (Figure 2). Therefore, we examined whether a difference in the leaf HPR activities between the two taxonomic groups could explain the observed difference in leaf glycerate contents. We found no significant difference in the mean HPR activity between the angiosperm trees and the conifers in the leaves collected in 2019 (Figure 4). Regarding the samples collected in 2020, the mean HPR activity of the conifer leaves was slightly lower than that of the angiosperm leaves. Thus, the activities of leaf HPR did not fully explain the lower glycerate levels detected in conifer leaves.

**Figure 4.**
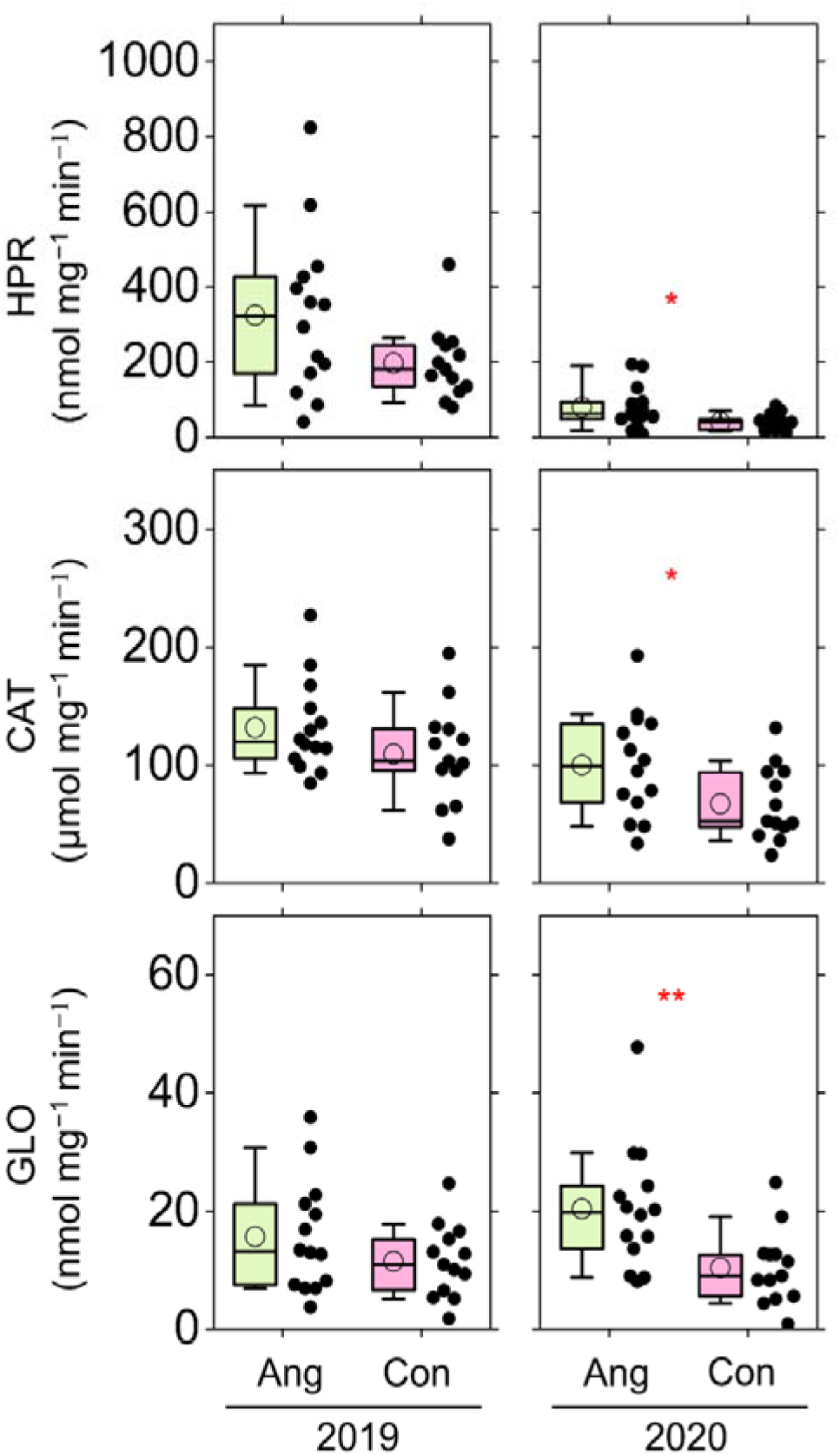
The activities of three enzymes, i.e., hydroxypyruvate reductase (HPR), catalase (CAT), and glycolatc oxidase (GLO), were determined in the leaves of 14 angiosperm tree (Ang) and those of 13 conifer (Con) species grown in a field site. The activities were expressed per unit soluble protein content. The left and right panels represent samples collected in 2019 and those collected in 2020, respectively. The central box in each box plot represents the interquartile range and median; the whiskers indicate the 10^th^ and 90^th^ percentiles. The open circles indicate the average values. The scatter plots represent the data distributions. Mann Whitney *U* tests were conducted for pairwise comparisons between conifers and angiosperms: *, P < 0.05; **, P < 0.01.

An in vitro experiment using isolated peroxisomes from the leaves of spinach beet (*Beta vulgaris*) suggested that OH-Pyr is decarboxylated via a non-enzymatic attack of H_2_O_2_, and is then converted to glycolate (Walton and Butt, 1981). Peroxisomes contain H_2_O_2_-generating and H_2_O_2_-scavenging enzymes, i.e., glycolate oxidase (GLO) and catalase (CAT), respectively. We predicted that conifer leaves would show a reduction in CAT activities, and that H_2_O_2_ would remain and react with OH-Pyr, thus explaining the lower glycerate and higher glycolate concentrations observed in the conifer leaves (Figure 1). Regarding the leaves collected in 2019, there were no significant difference in the CAT and GLO activities between angiosperms and conifers (Figure 4). In turn, in the samples collected in 2020, the mean CAT activity of the conifer leaves was slightly lower than that of the angiosperm leaves, whereas the activities of GLO and CAT were also decreased. Taken together, these results led us to conclude that there was not a specific reduction in CAT activities in the conifer leaves.

### Gene expression and amino acid structure of conifer catalases

The CAT and GLO proteins are peroxisomal targeting enzymes that physically interact with each other, thus contributing to the efficient scavenging of GLO-generated H_2_O_2_ in rice leaves (Zhang et al., 2016). This finding intrigued us to reconsider the cellular localization of these enzymes in conifers, first based on amino acid sequences.

Analyses of the cDNA libraries revealed that *P. nigra* and *C. japonica* carried three and two genes that encode CAT enzymes, respectively, which we termed *PnCAT1–3* for *P. nigra* and *CjCATA* and *CjCATB* for *C. japonica*. We confirmed the expression of these *CAT* genes in planta using qRT-PCR, the results of which indicated that the overall expression of the *CAT* genes was higher in the leaves than in the roots of *P. nigra*, whereas it was higher in the roots vs. the leaves of *C. japonica*. In *P. nigra*, the expression of *PnCAT2* was predominant among these three genes in the leaves (Figure 5, A). The expression of *PnCAT1* and *PnCAT2* was higher in the leaves than it was in the roots, whereas *PnCAT3* was almost equally expressed in the two organs. In *C. japonica*, *CjCATA* was expressed at a higher level in the leaves than in the roots, whereas the opposite pattern was detected for *CjCATB* (Figure 5, A).

**Figure 5.**
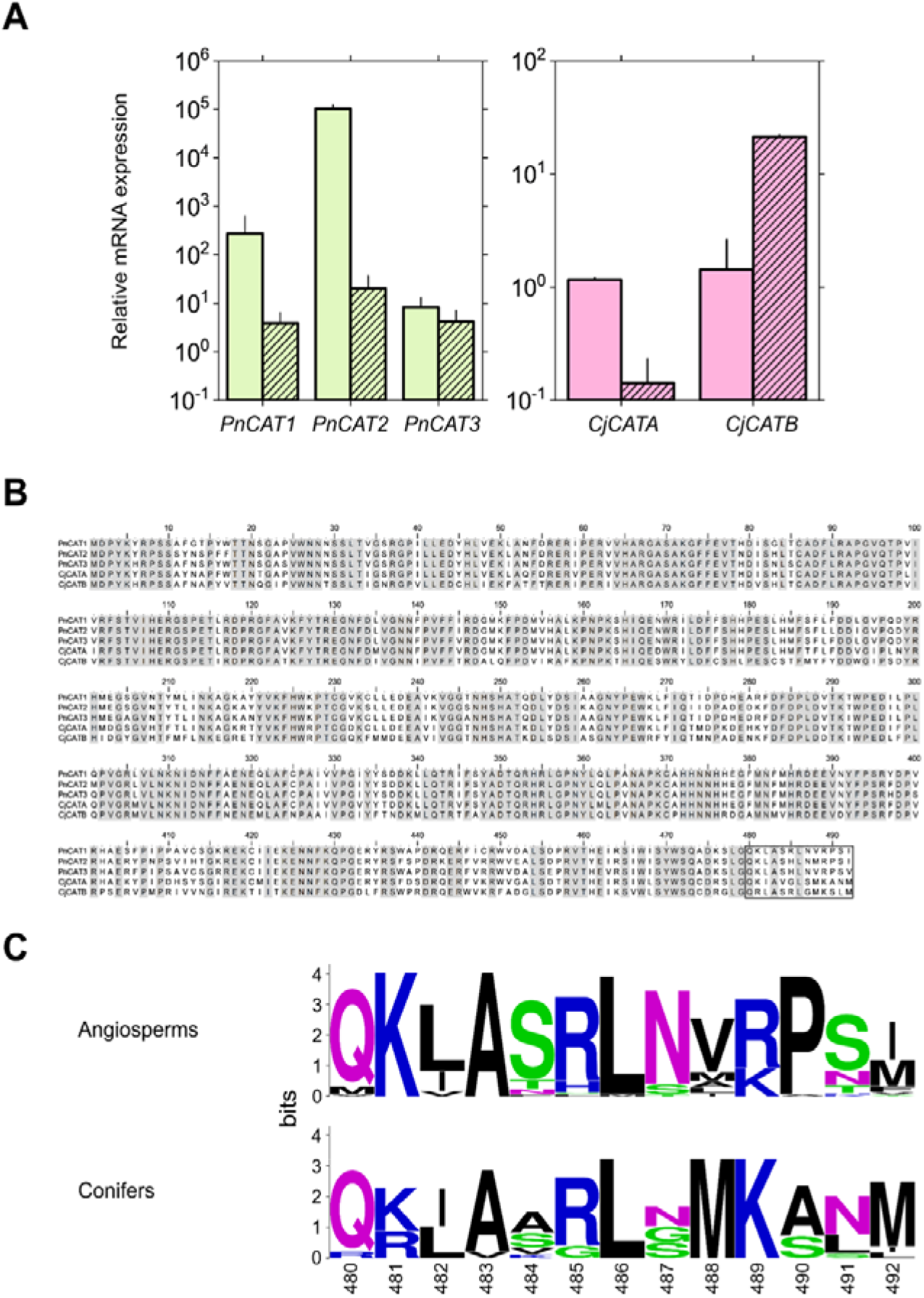
Differences in the gene expression and protein sequences of catalase (CAT) between angiosperms and conifers. A, Relative mRNΛexpression levels of the *CAT*genes in the leaves and roots of an angiosperm tree (*Populus nigra*) (left. *PnCATl-3*)and a conifer (*Cryptonicria japonica*) (right. *CjCATA* and *CjCATB*)saplings. The open and hatched bars represent the leaves and roots, respectively. The expression levels of the *CAT*genes were normalized against the expression level of the actin-encoding genes. Mean ±SD. *n*=3 biological replicates. B. Alignment of the amino acid sequences of the CAT proteins of *P. nigra* with those of C *japonic a*. Identical amino acid regions are indicated in gray. The region of the peroxisomal targeting signal (PTS), which has been implicated in the regulation of the import into peroxisomes, is boxed. C, Weblogo images of the PTS sequences of the CAT proteins for angiosperms and conifers. To generate the images, we referred to 51 genes from 18 angiosperm species, including herbs, grasses, and trees; and 13 genes from nine coniferous species, based on databases (Supplemental Table S3).

The 13 amino acids located at the carboxy-terminal (C-terminal) regions of CAT enzymes contain a peroxisomal targeting signal (PTS) motif, which was elucidated by in vivo subcellular targeting analyses, such as those using tobacco BY-2 cells and transgenic *Arabidopsis* plants (Mhamdi et al., 2010). The extreme C-terminal tripeptide is a sequence that is necessary for the efficient import of cottonseed CAT into BY-2 peroxisomes (designated as the P-S/T/N-M/I sequence by Mullen et al., 1997; Mhamdi et al., 2010).

We compared the deduced amino acid sequences of PnCATs and CjCATs, which exhibited overall high similarities between their CAT protein sequences, with the exception of a low similarity in the C-terminal regions (Figure 5, B). Moreover, we investigated the C-terminal sequences of CATs from 18 angiosperm species and nine coniferous species, and found that the Pro residue located at position –3 from the C terminus was well conserved among angiosperm CATs, including PnCAT1–3, with the exception of one of the maize CATs (ZmCAT3, NP_001350821.1) (Figure 5, B and C; Supplemental Table S3). In ZmCAT3, Pro was replaced with Ala. An organelle fractionation study indicated that ZmCAT3 was located in the mitochondrial fractions (Scandalios et al., 1980). Moreover, an analysis of the deduced amino acid sequences retrieved from several databases showed a clear difference in this amino acid between angiosperms and conifers: Pro was replaced with either Ala or Ser in all studied conifer CAT enzymes, including CjCATA and CjCATB (Figure 5, B and C). In contrast, the amino acids from positions −4 to −13 in the PTS did not show a specific difference between the angiosperm and the conifer CAT enzymes. Based on these results, we concluded that the conifer CAT enzymes were not localized at, or were less imported into, the peroxisomes.

Unlike CAT enzymes, an extreme C-terminal tripeptide functions as a PTS in GLO enzymes (Reumann et al., 2006). Based on the conifer database analyses, the C-terminal sequence of their GLOs exhibited the major PTS motifs, either S-K-L or S-R-L (Supplemental Table S4). Both motifs were experimentally verified by in vivo subcellular targeting analyses using transgenic *Arabidopsis* plants (Hayashi et al., 1997). Similarly, other photorespiration-related enzymes, including glutamate-glyoxylate aminotransferase (GGAT), HPR, malate dehydrogenase (MDH), and SGAT—which are the peroxisomal targeting enzymes in angiosperms—possessed the experimentally verified PTS motifs in all studied conifers (Supplemental Table S5).

### *Peroxisome fractions isolated from* Populus *and* Cryptomeria *leaves*

Our sequence analyses on the PTS motifs suggest that the conifer peroxisomes are not a major localization of CAT enzymes (Figure 5). To examine this finding experimentally, we isolated peroxisomes from *C. japonica* leaves using the Percoll/sucrose density gradient method. The GLOs of *C. japonica* and *P. nigra* had experimentally verified PTS motifs, i.e., S-R-L and P-R/K-L, respectively (Supplemental Table S4; Kaur et al. 2009), which allowed us to use GLO activity as a marker for elucidating peroxisome targeting. The measurement of glycerate kinase (GLYK) and fumarase (FUM) activities can be used to evaluate contaminations with chloroplasts and mitochondria in isolated peroxisome fractions (Usuda and Edwards, 1980). The calculated average recovery rates were 0.23% and 1.2% for *P. nigra* and *C. japonica*, respectively (Table 1), which were lower than the reported value for *Arabidopsis* (3.7%; Reumann et al., 2007). The calculated contamination with mitochondria (*C*_mito_.) and chloroplasts (*C*_chloro_.) was 16% and 7% for *C. japonica*, respectively (Figure 6). For *P. nigra*, *C*_mito_. was 3.7%, whereas significant *C*_chloro_. was not detected. As indicated by the peroxisomal localization index of CAT (*L*_CAT_), more than 90% of the leaf CAT was localized in peroxisomes in *P. nigra*,whereas only 13% of the leaf CAT was localized in peroxisomes in *C. japonica* (Figure 6). The low *L_CAT_* calculated for *C. japonica* was consistent with the results obtained for the PTS motif (Figure 5, C).

**Figure 6.**
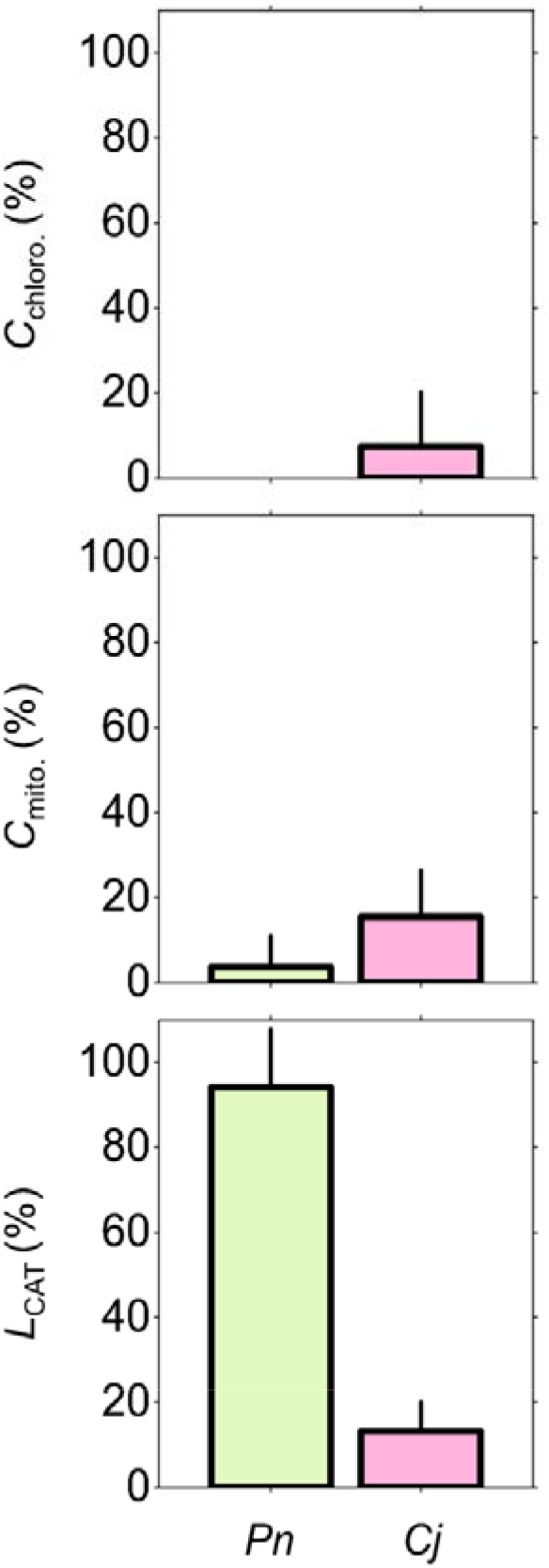
Characteristics of the peroxisome fractions isolated from the leaves of an angiosperm tree (*Populus nigra, Pn*) and from the leaves of a conifer (*Cryptomeria japonica, Cj*) using the Percoll/sucrose density gradient method. C_chloro_, chloroplast contamination index; C_mito_, mitochondrial contamination index; L_CAT_, peroxisomal localization index of catalase enzymes in the isolated peroxisome fractions. Mean ± SD from threc-to-four independent experiments (Table 1). The values of n.d. in Table 1 were assumed to be zero for the mean calculations.

### Chloroplastic CO_2_ compensation point (Γ*)

The *Γ*\* of *P. nigra* was unknown. Therefore, the *Γ*\* value of *P. nigra* leaves was estimated using the same method as that applied for *C. japonica* leaves (Miyazawa et al., 2020): *Γ*\* was estimated through measurements of the changes in the leaf CO_2_ compensation points (*Γ*) in response to the O_2_ partial pressure in air for a variety of leaves with different rates of Rubisco maximum carboxylation (*V*_cmax_) and respiration in the light (*R*_day_). The calculated *Γ*\* values (mean ± SE) were similar between the two species: 40.1 ± 0.9 and 41.2 ± 1.6 μbar for *P. nigra* and *C. japonica*, respectively (Supplemental Table S6; Supplemental Figure S2).

## Discussion

### The photorespiratory mechanism of conifers is different from that of angiosperm C_3_ species

The results of our metabolite and ^13^C-labeling analyses suggest that the photorespiratory mechanism is not identical between angiosperm tree and conifer leaves (Figure 2; Figure 3). Our findings from the analyses of the PTS motifs and isolated peroxisome fractions suggest that the H_2_O_2_-scavenging enzyme, CAT, is not abundant in conifer peroxisomes (Figure 5; Figure 6). Based on these findings, we propose a novel model for conifer photorespiration: a pathway from Ser to glycolate via the oxidative decarboxylation of OH-Pyr with H_2_O_2_, caused by an imbalance between CAT and GLO activities in peroxisomes. The present study was the first report showing that this decarboxylation pathway probably has a major flux in conifer photorespiration, and revealing that a diversification of photorespiratory metabolism exists among tree groups.

Previous studies have suggested that the oxidative decarboxylation of OH-Pyr occurs in transgenic tobacco plants, as well as in *Arabidopsis* mutants, which have impairments in peroxisome-targeting enzymes, such as CAT, HPR, and peroxisomal MDH (Brisson et al., 1998; Cousins et al., 2008, 2011). For example, transgenic tobacco with reduced CAT activities exhibited higher leaf CO_2_ compensation points (*Γ*) than the wild-type plants (Brisson et al., 1998). The *Γ* increased concomitantly with the decrease in the leaf CAT activities, probably because the excess of H_2_O_2_ caused by insufficient CAT activity decarboxylates ketoacids such as OH-Pyr and glyoxylate in peroxisomes, thereby causing a loss of assimilated CO_2_ (Brisson et al., 1998). This suggests that an imbalance between CAT and GLO activities in the peroxisomes caused an increase in the rate of photorespiratory CO_2_ release per unit *V_o_* (*α*). Conversely, the results pertaining to *Γ* warrant careful interpretation, because *Γ* is affected not only by the *α* values, but also by the *R*_day_/*V*_cmax_ ratio of the leaves, as noted in Equations 9 and 10 (Supplemental Methods).

The chloroplastic CO_2_ compensation point (*Γ*\*) is more simply defined than *Γ* because *Γ*\* is independent of *R*_day_/*V*_cmax_ (see Equation 3). The *Γ*\* values of *Arabidopsis* mutants lacking either peroxisomal MDH or HPR were about 1.5 times higher than those of the wild-type plants, because of the increases in *α* triggered by oxidative decarboxylation reactions of ketoacids in peroxisomes (Cousins et al., 2008, 2011). The present study suggests that such peroxisomal oxidative decarboxylation occurs in conifer leaves. In contrast, the *Γ*\* value of *C. japonica* leaves was similar to the values reported for the leaves of wild-type angiosperm species, such as *P. nigra* (this study), tobacco, and French bean (Miyazawa et al., 2020).

### The isolation process of conifer peroxisomes warrants improvement

The significantly low level of *L_CAT_* detected in *C. japonica* suggests that peroxisomes possess a lower amount of CAT compared with other subcellular fractions in conifers (Figure 6). Conversely, in the present study, we were not able to determine which subcellular fraction, i.e., mitochondria, chloroplasts, or the cytosol, etc., possessed a high level of CAT in the conifer leaf cells.

The peroxisome fractions of *Arabidopsis* were isolated with higher purity than ours: the organelle contaminations detected in *Arabidopsis* (*C*_chloro_. of 0.12% and *C*_mito_. of 1.7%; Reumann et al., 2007) were lower than those recorded in our studied species, particularly in *C. japonica* (Figure 6). Reumann et al. (2007) applied a two-step density gradient centrifugation method for the isolation of peroxisomes from *Arabidopsis* leaves: Percoll/sucrose density gradient followed by sucrose density gradient centrifugations. In our experiments, the Percoll/sucrose density gradient alone was applied for the isolation of peroxisomes. We attempted to isolate the peroxisomal fractions using the two-step method; however, we were not able to obtain measurable amounts of the peroxisomal fractions from the leaves of the two studied species. The low, but non-negligible, CAT activity detected in the isolated peroxisome fractions of *C. japonica* could be attributed to a relatively high contamination with chloroplasts and mitochondria.

Recently, in the leaves of rice, Zhang et al. (2016) reported that the CAT and GLO proteins associated together in peroxisomes and efficiently scavenged the H_2_O_2_ generated by GLO. In contrast, the conifer peroxisomes are not likely to possess such physical interaction between the CAT and GLO proteins, because the peroxisomes were not the major subcellular localization of CAT (Figure 6). However, we cannot rule out the possibility that other enzymes, such as ascorbate peroxidase, potentially scavenge H_2_O_2_ in peroxisomes (Kaur et al., 2009) and are involved in photorespiration in conifers. To elucidate more precisely the localization and functions of CAT in conifer leaf cells, we will need to improve the purity of the peroxisomes isolated from conifer leaves, and to perform additional subcellular fractionation analyses.

### Metabolites in the GDC and GS–GOGAT reactions

In the ^13^C-Ser feeding experiments, Gly was significantly labeled with ^13^C in the two studied species (Figure 3). Because glycolate was not significantly labeled with ^13^C in *P. nigra*, the significant labeling of Gly in this species might result from the synthesis of Gly from Ser via the reverse reaction of GDC in mitochondria (Sarojini and Oliver, 1983). It is also noteworthy that the concentrations of metabolites including Gln, 2-OG, and NH_4_^+^, which are involved in the GS–GOGAT cycle, were significantly different between the conifer and angiosperm tree leaves (Figure 2). The higher NH_4_^+^ content per unit chlorophyll concentration detected in the conifer leaves was in good agreement with the previous finding that conifer leaves have a higher NH_3_ emission potential than do angiosperm leaves (Miyazawa et al., 2018). Moreover, the concentration of 2-OG in the conifer leaves was only about one-tenth that of the angiosperms, whereas the Gln concentration in the conifer leaves was about two times higher (Figure 2). Although we do not have a straightforward elucidation of the physiological mechanisms that are responsible for such differences in the GS–GOGAT metabolites between the two taxonomic groups, it might be related to the fact that conifer leaves lack GS2 enzymes (Cánovas et al., 2007; Miyazawa et al., 2018).

### Inconsistency between Γ* and ^13^C-labeling experiment results

Our metabolite profile and ^13^C-labeing experiment results suggested that conifer leaves have an additional CO_2_ release from peroxisomes and, hence, a higher stoichiometry of photorespiratory CO_2_ release. Conversely, the in vivo *Γ*\* values of the conifer leaves were similar to those of the angiosperm leaves (Miyazawa et al., 2020). As noted in Equation 3, this inconsistency can be explained by three possibilities, assuming that the chloroplastic O_2_ concentration (*O*) is constant: (1) a higher *S*_c/o_, (2) a higher proportion of photorespiratory CO_2_ that is re-fixed by Rubisco, and (3) a lower rate of CO_2_ release by the GDC reactions per *V*_o_, in conifer leaves.

The three possibilities are explained in greater detail as follows. (1) Galmés et al. (2014) indicated that the CO_2_ affinity of Rubisco purified from the leaves of a conifer (*Metasequoia*) was significantly lower than that of Rubisco purified from the leaves of angiosperms. Therefore, it is unlikely that the *S_c/o_* of conifers is higher than that of angiosperms, although the kinetic data of conifer Rubisco is insufficient, and the in vitro *S*_c/o_ value for conifer Rubisco has not been reported to date. (2) The rate of CO_2_ release escaping from inside the leaves via the photorespiratory pathway is affected by the extent to which photorespiratory CO_2_ is re-fixed by Rubisco (Busch et al., 2013). The extent of this process is predicted to be larger in leaves with a thicker mesophyll cell wall that imposes a larger diffusion resistance to the CO_2_ efflux from inside the mesophyll cells to the intercellular air space in leaves (Tholen et al., 2012; Eckert et al., 2021). Because conifer leaves have thicker mesophyll cell walls than angiosperms, such as broad-leaved trees and C3 annual herbs (Terashima et al., 2011; Veromann-Jürgenson et al., 2017), the high stoichiometry of photorespiratory CO_2_ release in conifers can be cancelled out by a high rate of CO_2_ re-fixation by Rubisco. (3) The intermediates in the photorespiratory pathway, such as Gly and Ser, can be exported from the pathway and utilized in other metabolic processes, such as amino acid and secondary product syntheses, which could result in a change in the rate of CO_2_ release by the GDC reactions per *V*_o_

(Busch et al., 2020). Further studies are needed to investigate whether there are differences in the metabolic processes linked with the photorespiratory pathways between conifers and angiosperms, and whether the rate of CO_2_ release by GDC reactions per *V_o_* differs.

### Conclusions

The present study suggested that the differences in the subcellular localizations of CAT and GLO yielded a bypass from Ser to glycolate via the oxidative decarboxylation of OH-Pyr in conifer photorespiration. Together with the previous finding of an absence of GS2 in conifer leaves (Cánovas et al., 2007; Miyazawa et al., 2018), the present study supported the contention that conifer gymnosperms have a different photorespiratory mechanism compared with angiosperm C3 species. It is known that H_2_O_2_ decarboxylates not only OH-Pyr, but also glyoxylate, which is then converted to formate (Walton and Butt, 1981). We plan to investigate whether formate in leaves is also labeled by ^13^C after feeding ^13^C-Ser to detached shoots of *C. japonica*. The HPR requires the reductant NADH for converting OH-Pyr to glycerate (Ogren, 1984); in contrast, the oxidative decarboxylation may not have this requirement, implying that the energy requirements for the conifer photorespiratory pathway can be different from those in this pathway in angiosperms. Notably, our first observation of the differences in photorespiratory mechanisms between conifers and angiosperm C_3_ species suggests that the application of the FvCB model to conifer photosynthesis needs to be revised.

## Materials and Methods

### Plant materials

Analyses of the photorespiratory-metabolite profiling and enzymatic activity were performed using leaf samples collected from the current-year branches of the tree species listed in Supplemental Table S1. The tree species were cultivated in an FFPRI tree garden. Sunlit leaves were collected from three individuals per species (two individuals for *Morus australis*) between 10:00 and 14:00 on sunny days in August. The collected leaf samples were pooled for each species. The samplings were carried out in 2019 and 2020. For ^13^C-labeled Ser feeding and peroxisome-isolation experiments, we used potted saplings of *Cryptomeria japonica* and *Populus nigra*. Those plants were grown in an air-conditioned glasshouse under a natural light environment (day/night air temperature, 25°C/20°C; day/night relative humidity, 65%/60%) or under an artificial-light environment (day/night air temperature, 25°C/20°C; relative humidity, 70%; photosynthetically active photon flux density at the plant top, 400 μmol m^−2^ s^−1^). For gene expression analyses, the mature leaves of current-year branches and fine roots were sampled during the daytime (10:00–11:00) from three potted individuals for each species grown in the glasshouse in June for *P. nigra* and in September for *C. japonica*. The fine roots were cut with a pair of scissors and then rinsed with distilled water. The detailed growth conditions are described in the Supplemental Methods. All leaf and root samples were frozen in liquid nitrogen immediately after collection, and stored at −80°C until measurements.

### Metabolite determinations

Leaf metabolites were extracted using a chloroform and methanol solvent. The detailed procedures of extraction were as described in our previous paper (Tahara et al., 2021). Photorespiration-related organic acids other than glyoxylate were quantified using a GC-MS system (7890B/5977A; Agilent Technologies, Santa Clara, CA) equipped with a DB-5MS capillary column (30 m, 0.25 mm i.d., 0.25 μm film thickness, Agilent); whereas the amino acids were quantified using an HPLC system (Agilent 1100; Agilent) equipped with a Zorbax Eclipse Plus C18 column (4.6 mm × 150 mm, 3.5 μm particle size, Agilent).

To prepare GC-MS samples, the ketone moieties of the compounds in freeze-dried leaf extracts were methoximated using 10 μL of 40 mg mL^−1^ methoxiamine hydrochloride (FUJIFILM Wako Chemicals, Osaka, Japan) in pyridine at 30°C for 90 min; the acidic protons were subsequently derivatized with 90 μL of *N*-methyl-*N*-trimethylsilyltrifluoroacetamide plus 1% trimethylchlorosilane (Thermo Fisher Scientific, Waltham, MA) at 37°C for 30 min. Ribitol (final concentration, 4 μg mL^−1^) was used as an internal standard. The GC-MS conditions were exactly the those described by Tahara et al. (2021).

The GC-MS *m/z* values in selected ion monitoring mode were: 205, 217, 319 for ribitol; 205, 217, 292, 307 for glycerate; 147, 161, 177, 205 for glycolate; and 198, 229, 288, 304 for 2-OG. The GC-MS retention times (in min) were: 15.6 for ribitol; 10.7 for glycerate; 7.08 for glycolate; and 13.8 for 2-OG. Their amounts were determined based on their peak areas using authentic standards (Cat No. 1021-58400, GL Science, Tokyo, Japan). All calculations were carried out using the software (MassHunter, Quantitative Analysis Ver.7.01 SP1; Agilent). Glyoxylate was not within the measurable range of the GC-MS retention time. Therefore, glyoxylate concentrations were spectrophotometrically determined (UV-1900; Shimadzu, Kyoto, Japan) according to the method described by Häusler et al. (1996) with a slight modification. We used a 0.1 M HCl solution containing 1% (w/v) phenylhydrazine (instead of the 0.1% described in Häusler et al.).

The amino acid concentrations were determined by the HPLC method of pre-column derivatization using an *o*-phthalaldehyde and 9-fluorenylmethyl chloroformate reagent. Further details are described in the manufacturer manual (https://www.agilent.com/cs/library/applications/5990-4547EN.pdf, Agilent). Norvaline (final concentration, 8.4 μg mL^−1^) was used as an internal standard. Leaf chlorophyll was extracted with 80% acetone and spectrophotometrically quantified (Porra et al., 1989).

### Ammonium (NH_4_^+^)

Leaf NH_4_^+^ concentrations were fluorometrically determined using the *o*-phthalaldehyde (OPA) method in a neutral pH condition using a microplate reader (Varioscan LUX; Thermo Fisher Scientific), which was based on the methods described by Husted et al. (2000). A frozen leaf sample (20–40 mg of fresh weight per sample) was ground with 500 μL of 20 mM formate using a chilled mortar and pestle. Aliquots of leaf extracts (20 μL per sample) after centrifugation at 16000 × *g* (5 min at 4°C) were mixed with 180 μL of OPA solution in a 96-well plate placed on crushed ice. Subsequently, the plates were gently shaken at 60°C for 3 min (Microincubator M-36; TAITEC Co. Saitama, Japan) to proceed the reaction of NH_4_^+^ in extracts with OPA, and the florescence intensities at 470 nm (excited at 410 nm) were measured. The OPA solution contained 100 mM phosphate buffer (pH 7), 29 mM OPA, and 10 mM 2-mercaptoethanol. NH_4_NO_3_ was used as a standard.

### Leaf HPR, CAT, and GLO activities

For HPR activity measurements, frozen leaf samples were ground using a chilled mortar and pestle and homogenized in an extraction buffer containing 100 mM HEPES-KOH (pH 7.8), 1 mM EDTA, 10% (v/v) glycerol, 10 mM MgSO_4_, 5 mM dithiothreitol (DTT), 0.01 mM leupeptin, and 1% (v/v) Nonidet P-40. The supernatants of the extracts obtained after centrifugation were used for measurements. The HPR activity was determined based on the decrease in absorbance at 340 nm in an assay solution containing 100 mM HEPES-KOH (pH 6.8), 0.1 mM NADH, and 1 mM lithium hydroxypyruvate (Husic and Tolbert, 1987) at a controlled temperature of 30°C. For CAT and GLO activity measurements, frozen leaf samples were ground using a chilled mortar and pestle and homogenized in an extraction buffer containing 50 mM phosphate buffer (pH 7.0), 1 mM EDTA, 10% (v/v) glycerol, 0.01 mM leupeptin, and 1% (v/v) Nonidet P-40. The supernatants of the extract obtained after centrifugation were used for measurements. CAT activity was determined using an O_2_ electrode (Oxygraph+; Hansatech Instruments Ltd., Norfolk, UK) (del Río et al., 1977). The final concentration of H_2_O_2_ used as a substrate was ca. 180 mM. GLO activity was spectrophotometrically determined based on the method described by Yamaguchi and Nishimura (2000). The measurement temperatures were controlled at 30°C. Further details are described in the Supplemental Methods. The soluble protein content was spectrophotometrically determined using bovine serum albumin as a standard (Bradford Protein Assay Kit; Bio-Rad, Hercules, CA).

### ^13^C-labeled Ser feeding

Before plants were illuminated, the shoots were enclosed with aluminum foil to keep them under dark conditions until the onset of the treatments. The shoots were detached from the plants using a razor blade and were immediately immersed in distilled water in 50-mL tubes (one shoot per tube). The shoots kept in the tubes were placed under a light intensity of 400 μmol m^−2^ s^−1^ in an air-conditioned growth chamber (25°C and 70% relative humidity) at an ambient CO_2_ concentration of ca. 400 μmol mol^−1^ for 30 min. Subsequently, labeled L-Ser, the C3-position of which is mainly labeled with a stable isotope of carbon (^13^C-Ser), was added to the tubes (final concentration, 1.6 or 3.2 mM). The distribution of the isotopologue abundance of an unlabeled Ser standard (uSer) was similar to that detected in the leaves sampled before ^13^C-Ser addition (Supplemental Figure S3). Therefore, uSer at the same concentrations as those of the corresponding ^13^C-Ser treatments was used as controls. ^13^C-Ser (Cat. No. 604720) and uSer (Cat. No. 54763) were purchased from MilliporeSigma (Burlington, MA). The shoots were sampled 20 or 50 min after Ser additions.

### Relative isotopologue abundance

The relative isotopologue abundance (*m*_i_) of each metabolite into which *i*^13^C atoms were incorporated was calculated using the following equation (Hasunuma et al., 2010):

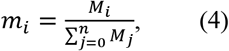

where *M_i_* represents the abundance of the isotopologues into which *i*^13^C atoms were incorporated and *n* is the number of C atoms that each metabolite possessed. According to Berry et al. (1978) and Huege et al. (2007), *M*_i_ at the selected GC-MS *m/z* values was measured in the single-ion monitoring mode (Supplemental Table S7). The major isotopologue of ^13^C-Ser possessed a single ^13^C atom (*m*_1_ = 77%; Supplemental Figure S3). Therefore, a ^13^C enrichment of each photorespiratory metabolite was evaluated as a percent increase in *m*_1_ for the leaves that were fed with ^13^C-Ser from that recorded for the leaves that were fed with uSer.

### Peroxisome isolation

Leaf peroxisomes were isolated using the Percoll/sucrose density gradient method (Reumann and Lisik, 2017), with a slight modification. Our extraction buffer contained no DTT because severe inhibitions of CAT activities by DTT have been reported (Chandlee et al., 1983). The detached leaves enclosed in a plastic bag were kept on crushed ice in the dark (2 h), for loosening the physical interactions among organelles. The leaves were ground using a mortar and pestle on ice with an extraction buffer containing 170 mM Tricine-KOH (pH 7.5), 1.0 M sucrose, 2 mM EDTA, 1% (w/v) BSA, 10 mM KCl, 1 mM MgCl_2_, and 0.5% (w/v) polyvinylpyrrolidone-40 supplemented with a protease inhibitor cocktail (cOmplete, EDTA-free; Roche, Basel, Switzerland). After the removal of cellular debris, chloroplasts, and nuclei from the extracted solution via filtration using a layer of Miracloth (Millipore, MA) and mild centrifugation (5,000 × *g* for 1 min), 15 mL of the supernatant was loaded onto a Percoll/sucrose density gradient prepared in TE buffer (20 mM Tricine-KOH (pH 7.5) and 1 mM EDTA) supplemented with 0.75 M sucrose and 0.2% (w/v) BSA, and underlaid by 36% (w/v) sucrose in TE buffer (top to bottom: 2.4 mL of 15% Percoll, 8.8 mL of 38% Percoll, 1.6 mL of a mixture of 38% Percoll and 36% [w/v] sucrose at a ratio of 2:1 and 1:2, and 2.4 mL of 36% [w/v] sucrose in TE buffer). A sediment fraction corresponding to crude peroxisomal fractions (LP-P1 fraction; Reumann and Lisik, 2017) was obtained by stepwise centrifugation (13,000 × *g* for 12 min, followed by an increase to 27,000 × *g* for an additional 20 min at 4°C) on an ultracentrifuge (Himac SCP85H2; Hitachi, Tokyo, Japan). The isolated peroxisome fractions were stored in a freezer at −80°C until enzyme activity measurements. The CAT and GLO activities were measured as described above. The activities of GLYK and FUM were measured and used as markers for identifying chloroplast and mitochondria contaminations, respectively, at a temperature of 30°C (Kleczkowski and Randall, 1986; Gibon et al., 2004). The contamination of leaf peroxisomes by chloroplasts and mitochondria was calculated (Reumann et al., 2007) based on the relative activities of the corresponding marker enzymes (i.e., *Act*_GLYK_ and *Act*_FUM_) that were recovered in the leaf peroxisome (LP) fraction compared with the crude extract (CE). The activity of GLO (*Act*_GLO_) was used as a peroxisome marker. For example, the contamination with chloroplasts (*C*_chloro_.) was calculated as:

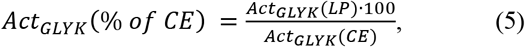

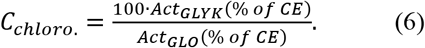

Using the same analogy of the contamination calculations, the peroxisome localization of CAT (*L*_CAT_) was calculated as:

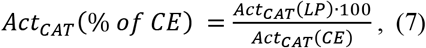

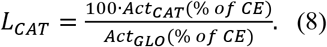

### Estimation of Γ* for Populus nigra

We estimated the *Γ*\* of *P. nigra* from the analyses of the O_2_ responses of the leaf CO_2_ compensation points (*Γ*) according to the method described by Miyazawa et al. (2020), and compared it with the reported *Γ*\* value of *C. japonica*. The initial slopes of the relationships between the leaf net CO_2_ assimilation rates (*A*) and the intercellular CO_2_ concentrations (*C*_i_) were obtained at different air-O_2_ concentrations using a portable CO_2_ gas exchange analyzer (LI-6400; Li-Cor, Lincoln, NE). The O_2_ concentrations ranged from 10% to 30% (10%, 15%, 20%, 25% and 30%). The *A*-*C*_i_ initial slope was fitted by linear regression. The value at the location on the *x*-axis where the linear regression lines crossed was taken as the estimate of *Γ*. Leaf temperature was controlled at 25°C. For additional details on the measurement procedures and conditions, refer to the Supplemental Methods.

### Sequencing

The catalase-encoding genes of *P. nigra* and *C. japonica* were searched against cDNA libraries (ForestGEN; http://forestgen.ffpri.affrc.go.jp/en/index.html) using an *Arabidopsis* catalase sequence (GenBank No.; NP_564121.1) as a query on the TBLASN program. The selected cDNA clones were sequenced on a 3130xl Genetic Analyzer (Applied Biosystems, Foster City, CA). The deduced amino acid sequences were aligned using the MUSCLE program (GENETYX ver. 15.0.0; Genetyx, Tokyo, Japan). The sequence logos of the peroxisomal targeting regions were generated using Weblogo (https://weblogo.berkeley.edu/logo.cgi).

### Quantitative real-time RT-PCR (qRT-PCR)

Frozen leaf samples were ground using a chilled mortar and pestle. Total RNA was isolated according to the method described by Izuno et al. (2020), and subsequently used for the synthesis of the first-strand cDNA with iScript Reverse Transcription Supermix (Bio-Rad) after checking the RNA quality (2100 Bioanalyzer; Agilent). The quantitative real-time PCR assay was performed using the SsoAdvanced SYBR Green Supermix (Bio-Rad) with specific primers (Supplemental Table S8). The expression levels of the *CAT* genes were normalized to the expression level of the respective actin-encoding genes (*PnACT* or *CjACT*).

### Statistical analysis

Mann–Whiney *U* tests and Student’s *t*-tests were performed using the Stata 16.1 software (StataCorp, College Station, TX). The box plots were generated using a graphing computer software (Origin 2021b; Origin Lab., Northampton, MA).

## Accession Numbers

Genes referenced in this article are available at GenBank with the accession numbers: LC627279 (*PnCAT1*), LC627280 (*PnCAT2*), LC627281 (*PnCAT3*), LC627277 (*CjCATA*),LC627278 (*CjCATB*).

## Supplemental Data

The following materials are available in the online version of this article.

Supplemental Methods.

Supplemental Figure S1. Metabolite concentrations on a leaf fresh weight basis.

Supplemental Figure S2. Relationships between *d* and *R*_day_/*V*_cmax_.

Supplemental Figure S3. Isotopologue abundance of Ser.

Supplemental Table S1. Photorespiratory-metabolite concentrations.

Supplemental Table S2. Isotopologue abundance of the ^13^C-Ser feedings.

Supplemental Table S3. PTS motifs of catalase (CAT).

Supplemental Table S4. PTS motifs of glycolate oxidase (GLO).

Supplemental Table S5. PTS motifs of photorespiratory enzymes other than CAT and GLO.

Supplemental Table S6. Gas exchange parameters of *P. nigra* leaves.

Supplemental Table S7. *m/z* values for the ^13^C-Ser feedings.

Supplemental Table S8. List of the primers used in the CAT expression analyses.

## Acknowledgements

We are grateful to Ms. Rie Yamamoto in FFPRI for her kind help in running the experiments, and also Prof. Ko Noguchi in Tokyo University of Pharmacy and Medical Sciences for his constructive comments on this research project.

## Parsed Citations

Berry JA, Osmond B, Lorimer GH (1978) Fixation of 18O2 during photorespiration: Kinetic and steady-state studies of the photorespiratory carbon oxidation cycle with intact leaves and isolated chloroplasts of C3 plants. Plant Physiol 62: 954–967 Google Scholar:Author Only Title Only Author and Title

Brisson LF, Zelitch I, Havir EA (1998) Manipulation of catalase levels produces altered photosynthesis in transgenic tobacco plants. Plant Physiol 116: 259–269 Google Scholar: Author Only Title Only Author and Title

Busch FA, Sage TL, Cousins AB, Sage RF (2013) C3 plants enhance rates of photosynthesis by reassimilating photorespired and respired CO2. Plant Cell Environ 36: 200–212 Google Scholar: Author Only Title Only Author and Title

Busch FA (2020) Photorespiration in the context of Rubisco biochemistry, CO2 diffusion and metabolism. Plant J 101: 919–939 Google Scholar: Author Only Title Only Author and Title

Cánovas FM, Avila C, Cantón FR, Cañas RA, de la Torre F (2007) Ammonium assimilation and amino acid metabolism in conifers. J Exp Bot 58: 2307–2318 Google Scholar: Author Only Title Only Author and Title

Chandlee JM, Tsaftaris AS, Scandalios JG (1983) Purification and partial characterization of three genetically defined catalases of maize. Plant Sci Lett 29: 117–131 Google Scholar: Author Only Title Only Author and Title

Cousins AB, Pracharoenwattana I, Zhou W, Smith SM, Badger MR (2008) Peroxisomal malate dehydrogenase is not essential for photorespiration in Arabidopsis but its absence causes an increase in the stoichiometry of photorespiratory CO2 release. Plant Physiol 148: 786–795 Google Scholar: Author Only Title Only Author and Title

Cousins AB, Walker BJ, Pracharoenwattana I, Smith SM, Badger MR (2011) Peroxisomal hydroxypyruvate reductase is not essential for photorespiration in Arabidopsis but its absence causes an increase in the stoichiometry of photorespiratory CO2 release. Photosynth Res 108: 91–100 Google Scholar: Author Only Title Only Author and Title

del Río LA, Ortega MG, López AL, Gorgé JL (1977) A more sensitive modification of the catalase assay with the Clark oxygen electrode: application to the kinetic study of the pea leaf enzyme. Anal Biochem 80: 409–415 Google Scholar: Author Only Title Only Author and Title

Eckert D, Martens HJ, Gu L, Jensen AM (2021) CO2 refixation is higher in leaves of woody species with high mesophyll and stomatal resistances to CO2 diffusion. Tree Physiol 41: 1450–1461 Google Scholar: Author Only Title Only Author and Title

Farquhar GD, von Caemmerer S, Berry JA (1980) A biochemical model of photosynthetic CO2 assimilation in leaves of C3 species. Planta 149: 78–90 Google Scholar: Author Only Title Only Author and Title

Galmés J, Kapralov MV, Andralojc PJ, Conesa MÀ, Keys AJ, Parry MAJ, Flexas J (2014) Expanding knowledge of the Rubisco kinetics variability in plant species: environmental and evolutionary trends. Plant Cell Environ 37: 1989–2001 Google Scholar: Author Only Title Only Author and Title

Gibon Y, Blaesing OE, Hannemann J, Carillo P, Höhne M, Hendriks JHM, Palacios N, Cross J, Selbig J, Stitt M (2004) A robot-based platform to measure multiple enzyme activities in Arabidopsis using a set of cycling assays: comparison of changes of enzyme activities and transcript levels during diurnal cycles and in prolonged darkness. Plant Cell 16: 3304–3325 Google Scholar: Author Only Title Only Author and Title

Häusler RE, Bailey KJ, Lea PJ, Leegood RC (1996) Control of photosynthesis in barley mutants with reduced activities of glutamine synthetase and glutamate synthase. III. Aspects of glyoxylate metabolism and effects of glyoxylate on the activation state of ribulose-l,5-bisphosphate carboxylase-oxygenase. Planta 200: 388–396 Google Scholar: Author Only Title Only Author and Title

Hasunuma T, Harada K, Miyazawa S-I, Kondo A, Fukusaki E, Miyake C (2010) Metabolic turnover analysis by a combination of in vivo 13C-labelling from 13CO2 and metabolic profiling with CE-MS/MS reveals rate-limiting steps of the C3 photosynthetic pathway in Nicotiana tabacum leaves. J Exp Bot 61: 1041–1051 Google Scholar: Author Only Title Only Author and Title

Hayashi M, Aoki M, Kondo M, Nishimura M (1997) Changes in targeting efficiencies of proteins to plant microbodies caused by amino acid substitutions in the carboxy-terminal tripeptide. Plant Cell Physiol 38: 759–768 Google Scholar: Author Only Title Only Author and Title

Hermida-Carrera C, Kapralov MV, Galmés J (2016) Rubisco catalytic properties and temperature response in crops. Plant Physiol 171: 2549–2561 Google Scholar: Author Only Title Only Author and Title

Huege J, Sulpice R, Gibon Y, Lisec J, Koehl K, Kopka J (2007) GC-EI-TOF-MS analysis of in vivo carbon-partitioning into soluble metabolite pools of higher plants by monitoring isotope dilution after 13CO2 labelling. Phytochem 68: 2258–2272 Google Scholar: Author Only Title Only Author and Title

Husic DW, Tolbert NE (1987) NADH:hydroxypyruvate reductase and NADPH:glyoxylate reductase in algae: partial purification and characterization from Chlamydomonas reinhardtii. Arch Biochem Biophys 252: 396–408 Google Scholar: Author Only Title Only Author and Title

Husted S, Hebbern CA, Mattsson M, Schjoerring JK (2000) A critical experimental evaluation of methods for determination of NH4+ in plant tissue, xylem sap and apoplastic fluid. Physiol Plant 109: 167–179 Google Scholar: Author Only Title Only Author and Title

Izuno A, Maruyama TE, Ueno S, Ujino-Ihara T, Moriguchi Y (2020) Genotype and transcriptome effects on somatic embryogenesis in Cryptomeria japonica. PLoS ONE 15: e0244634 Google Scholar: Author Only Title Only Author and Title

Kaur N, Reumann S, Hu J (2009) Peroxisome biogenesis and function. Arabidopsis Book 7: e0123 Google Scholar: Author Only Title Only Author and Title

Kleczkowski LA, Randall DD (1986) Thiol-dependent regulation of glycerate metabolism in leaf extracts: the role of glycerate kinase in C4 plants. Plant Physiol 81: 656–662 Google Scholar: Author Only Title Only Author and Title

Mhamdi A, Queval G, Chaouch S, Vanderauwera S, Van Breusegem F, Noctor G (2010) Catalase function in plants: a focus on Arabidopsis mutants as stress-mimic models. J Exp Bot 61: 4197–4220 Google Scholar: Author Only Title Only Author and Title

Miyazawa S-I, Nishiguchi M, Futamura N, Yukawa T, Miyao M, Maruyama TE, Kawahara T (2018) Low assimilation efficiency of photorespiratory ammonia in conifer leaves. J Plant Res 131: 789–802 Google Scholar: Author Only Title Only Author and Title

Miyazawa S-I, Tobita H, Ujino-Ihara T, Suzuki Y (2020) Oxygen response of leaf CO2 compensation points used to determine Rubisco specificity factors of gymnosperm species. J Plant Res 133: 205–215 Google Scholar: Author Only Title Only Author and Title

Mullen RT, Lee MS, Trelease RN (1997) Identification of the peroxisomal targeting signal for cottonseed catalase. Plant J 12: 313–322 Google Scholar: Author Only Title Only Author and Title

Ogren WL (1984) Photorespiration: pathways, regulation, and modification. Ann Rev Plant Physiol 35: 415–442 Google Scholar: Author Only Title Only Author and Title

Porra RJ, Thompson WA, Kriedemann PE (1989) Determination of accurate extinction coefficients and simultaneous equations for assaying chlorophylls a and b extracted with four different solvents: verification of the concentration of chlorophyll standards by atomic absorption spectroscopy. Biochim Biophys Acta 975: 384–394 Google Scholar: Author Only Title Only Author and Title

Reumann S, Babujee L, Ma C, Wienkoop S, Siemsen T, Antonicelli GE, Rasche N, Lüder F, Weckwerth W, Jahn O (2007) Proteome analysis of Arabidopsis leaf peroxisomes reveals novel targeting peptides, metabolic pathways, and defense mechanisms. Plant Cell 19: 3170–3193 Google Scholar: Author Only Title Only Author and Title

Reumann S, Lisik P (2017) Isolation of Arabidopsis leaf peroxisomes and the peroxisomal membrane. In NL Taylor, AH Miller, eds, Isolation of Plant Organelles and Structures: Methods and Protocols. Springer, New York, pp 97–112 Google Scholar: Author Only Title Only Author and Title

Reumann S, Weber APM (2006) Plant peroxisomes respire in the light: Some gaps of the photorespiratory C2 cycle have become filled-Others remain. Biochim Biophys Acta 1763: 1496–1510 Google Scholar: Author Only Title Only Author and Title

Sarojini G, Oliver DJ (1983) Extraction and partial characterization of the glycine decarboxylase multienzyme complex from pea leaf mitochondria. Plant Physiol 72: 194–199 Google Scholar: Author Only Title Only Author and Title

Scandalios JG, Tong W-F, Roupakias DG (1980) Cat3, a third gene locus coding for a tissue-specific catalase in maize: genetics, intracellular location, and some biochemical properties. Mol Gen Genet 179: 33–41 Google Scholar: Author Only Title Only Author and Title

Somerville CR (2001) An early Arabidopsis demonstration. Resolving a few Issues concerning photorespiration. Plant Physiol 125: 20–24 Google Scholar: Author Only Title Only Author and Title

Suárez MF, Avila C, Gallardo F, Cantón FR, García-Gutiérrez A, Claros MG, Cánovas FM (2002) Molecular and enzymatic analysis of ammonium assimilation in woody plants. J Exp Bot 53: 891–904 Google Scholar: Author Only Title Only Author and Title

Tahara K, Nishiguchi M, Funke E, Miyazawa S-I, Miyama T, Milkowski C (2021) Dehydroquinate dehydratase/shikimate dehydrogenases involved in gallate biosynthesis of the aluminum-tolerant tree species Eucalyptus camaldulensis. Planta 253: 3 Google Scholar: Author Only Title Only Author and Title

Terashima I, Hanba YT, Tholen D, Niinemets Ü (2011) Leaf functional anatomy in relation to photosynthesis. Plant Physiol 155: 108–116 Google Scholar: Author Only Title Only Author and Title

Tholen D, Ethier G, Genty B, Pepin S, Zhu X-G (2012) Variable mesophyll conductance revisited: theoretical background and experimental implications. Plant Cell Environ 35: 2087–2103 Google Scholar: Author Only Title Only Author and Title

Usuda H, Edwards GE (1980) Localization of glycerate kinase and some enzymes for sucrose synthesis in C3 and C4 plants. Plant Physiol 65: 1017–1022 Google Scholar: Author Only Title Only Author and Title

Veromann-Jürgenson L-L, Tosens T, Laanisto L, Niinemets Ü (2017) Extremely thick cell walls and low mesophyll conductance: welcome to the world of ancient living! J Exp Bot 68: 1639–1653

Wallsgrove RM, Turner JC, Hall NP, Kendall AC, Bright SWJ (1987) Barley mutants lacking chloroplast glutamine synthetase-biochemical and genetic analysis. Plant Physiol 83: 155–158 Google Scholar: Author Only Title Only Author and Title

Walton NJ, Butt VS (1981) Metabolism and decarboxylation of glycollate and serine in leaf peroxisomes. Planta 153: 225–231 Google Scholar: Author Only Title Only Author and Title

Yamaguchi K, Nishimura M (2000) Reduction to below threshold levels of glycolate oxidase activities in transgenic tobacco enhances photoinhibition during irradiation. Plant Cell Physiol 41: 1397–1406 Google Scholar: Author Only Title Only Author and Title

Zhang Z, Xu Y, Xie Z, Li X, He Z-H, Peng X-X (2016) Association-dissociation of glycolate oxidase with catalase in rice: a potential switch to modulate intracellular H2O2 levels. Mol Plant 9: 737–748 Google Scholar: Author Only Title Only Author and Title

